# Deletion of carboxypeptidase E in beta cells disrupts proinsulin processing and alters beta cell identity in mice

**DOI:** 10.1101/2022.10.20.512925

**Authors:** Yi-Chun Chen, Austin J. Taylor, James M. Fulcher, Adam C. Swensen, Xiao-Qing Dai, Mitsuhiro Komba, Kenzie L.C. Wrightson, Kenny Fok, Annette E. Patterson, Ramon I. Klein-Geltink, Patrick E. MacDonald, Wei-Jun Qian, C. Bruce Verchere

## Abstract

Carboxypeptidase E (CPE) facilitates the conversion of prohormones into mature hormones and is highly expressed in multiple neuroendocrine tissues. Carriers of *CPE* mutations have elevated plasma proinsulin and develop severe obesity and hyperglycemia. We aimed to determine whether loss of *Cpe* in pancreatic beta cells disrupts proinsulin processing and accelerates development of diabetes and obesity in mice. Pancreatic beta cell-specific Cpe knockout mice (β*Cpe*KO; *Cpe*^fl/fl^ x *Ins1*^Cre/+^) lack mature insulin granules and have elevated proinsulin in plasma; however, glucose-and KCl-stimulated insulin secretion in β*Cpe*KO islets remained intact. High fat diet-fed β*Cpe*KO mice showed comparable weight gain and glucose tolerance compared to *Wt* littermates. Notably, beta-cell area was increased in chow-fed β*Cpe*KO mice and beta-cell replication was elevated in β*Cpe*KO islets. Transcriptomic analysis of β*Cpe*KO beta cells revealed elevated glycolysis and *Hif1α*-target gene expression. Upon high glucose challenge, beta cells from β*Cpe*KO mice showed reduced mitochondrial membrane potential, increased reactive oxygen species, reduced *MafA*, and elevated *Aldh1a3* transcript levels. Following multiple low-dose streptozotocin treatment, β*Cpe*KO mice had accelerated hyperglycemia with reduced beta-cell insulin and Glut2 expression. These findings suggest that *Cpe* and proper proinsulin processing are critical in maintaining beta cell function during the development of diabetes.

## Introduction

Proinsulin is processed into mature insulin and C-peptide by prohormone convertase 1/3 and 2 (PC1/3 and PC2) and carboxypeptidase E (CPE) (1–3) prior to secretion from pancreatic beta cells. Failure of this process leads to insufficient mature insulin release and onset of hyperglycemia (4–6) and has been observed in diabetes pathogenesis (7,8).

Prohormone processing enzymes are highly expressed in neuroendocrine cells and subjects with mutations in these genes often display cognitive impairments and obesity. A *CPE* truncating mutation (c.76_98del) causes morbid obesity and severe hyperglycemia (9), a *CPE* non-conservative missense mutation (c.847C>T) reduces enzymatic activity associated with early-onset type 2 diabetes (10), and homozygous nonsense *CPE* mutations (c.405C>A) cause obesity and hypogonadotropic hypogonadism (11). Similarly, *Cpe* whole-body knockout mice develop spontaneous obesity and behavioral abnormalities (12), and *Cpe* mutant mice (*Cpe*^fat/fat^, Ser202Pro mutation) are obese and infertile (13).

To understand the role of *Cpe* in beta-cell function and glucose homeostasis, we generated pancreatic beta-cell-specific *Cpe* knockout mice. We performed biochemical and top-down proteomic analysis to evaluate hormone processing patterns in beta cells. We also analyzed beta-cell transcriptomic profiles and performed live-cell imaging analysis to understand whether deficiency of *Cpe*, and the increased compensatory production of proinsulin, leads to islet dysfunction and dysglycemia. Finally, we tested whether the lack of *Cpe* in beta cells increases susceptibility to diet- or secretory stress-induced hyperglycemia in mice. Our model provides a useful tool to understand the role of reduced prohormone processing efficiency and increased (pro)insulin translation in beta cells during diabetes development, and sheds light on whether impaired prohormone processing in beta cells is a cause of diabetes and obesity in subjects with *CPE* mutations.

## Research Design and Methods

### Human Pancreas Tissue

Paraffin-embedded pancreatic tissue sections were obtained from the Network for Pancreatic Organ Donors with Diabetes (nPOD) and Alberta Diabetes Institute IsletCore (Supplementary Table 1).

### Mouse Studies

Beta-cell-specific *Cpe* knockout (β*Cpe*KO) and inducible beta-cell-specific *Cpe* knockout (iβ*Cpe*KO) mice were generated by crossing the offspring of C57BL/6N-Cpe^tm1a(EUCOMM)Hmgu^/Ieg mice and Tg(CAG-flpo)1Afst mice, with B6(Cg)-Ins1^tm1.1(cre)Thor^/J or Tg(Pdx1-cre/Esr1) mice. In addition, *Gt(ROSA)26Sor*^*tm4(ACTB-tdTomato,-EGFP)Luo*^ reporter mice were bred with β*Cpe*KO mice for beta-cell sorting and RNA sequencing. For diet studies, 8-week-old mice received either a low-fat diet (LFD; 10% fat), or a high-fat diet (HFD; 45% fat, Research Diets). For secretory stress studies, 10-week-old male mice received saline or multiple low-dose STZ (MLD-STZ; Sigma-Aldrich; 35 mg/kg body weight, daily i.p. for 5 days) injections. Metabolic assays (such as intraperitoneal glucose tolerance test (IPGTT), insulin tolerance test (ITT), and body mass composition analysis) were performed in a blinded fashion and have been described previously (14). For *in vivo* beta-cell proliferation studies, after tamoxifen-induced *Cpe* deletion, a 60% fat diet (Research Diet) was given for 48 hours (15) in combination with 5-ethynyl-2’-deoxyuridine (EdU; Toronto Research Chemicals, 40 mg/kg body weight, twice daily i.p. injection) (16). All studies were approved by the Animal Care and Use Committee at the University of British Columbia.

### Islet Studies

Mouse islets were isolated and cultured as described (14). For electron micrograph studies, freshly isolated islets were fixed in 2% glutaraldehyde (pH 7.4) at room temperature, shipped, processed, and imaged by the Electron Microscopy Facility at McMaster University Health Science Centre. PC1/3 and PC2 enzyme activity assays were performed as described (17) using a SpectraMax M3 plate reader (Molecular Devices). For respirometry studies, mouse islets were dispersed and analyzed by Seahorse XFe96 Analyzer (Agilent). To analyze insulin secretion dynamics, islets were incubated in a perifusion system (BioRep) with 1.67 mM glucose, 16.7 mM glucose, and 16.7mM glucose plus 30mM KCl Krebs-Ringer buffer (KRB) sequentially. Insulin concentrations in perifusates were analyzed by rodent insulin (Alpco) and proinsulin (Mercodia) ELISAs. To measure glucose uptake, islets were pre-cultured in glucose-free media, dispersed with Accutase (Innovative Cell), and treated with 2-NBDG (2-Deoxy-2-[(7-nitro-2,1,3-benzoxadiazol-4-yl)amino]-D-glucose; Invitrogen) for 5 minutes, prior to flow cytometry analysis.

For exocytosis studies, dispersed islet beta cells were patch-clamped in whole-cell voltage-clamp configuration using a HEKA EPC10 amplifier and PatchMaster Software (Heka Electronik, Germany) as described previously (18). Exocytosis was monitored as increases in cell capacitance, elicited by either a series of 500-ms membrane depolarizations from -70 to 0 mV or increasing the duration of membrane depolarizations. For FACS-sorted beta-cell bulk-RNA sequencing experiments, freshly isolated islets were dispersed and GFP^+^ live cells were collected by BD FACSAria™ Cell Sorter for RNA isolation via a RNeasy Plus Micro Kit (Qiagen). After quality control analysis using an Agilent 2100 Bioanalyzer, an RNA library was prepared using the NeoPrep Library Prep System with TruSeq Stranded mRNA Kit (Illumina), RNA sequencing was performed using Ilumina NextSeq500, reads aligned by TopHat to the reference genome of UCSC mm10, assembled by Cufflinks, and a list of differentially expressed genes were generated via DESeq2. Gene-set enrichment analysis, network visualization, and volcano plot were generated via Gene Set Enrichment Analysis (GSEA v4.2.3), Cytoscape (v3.9.1), and EnhancedVolcano (v1.14.0) in RStudio (v1.4.1717).

### Top-down Proteomic Analysis

Islet pellets were homogenized in 8 M urea lysis buffer, reduced, alkylated, quenched, and clarified with tris(2-carboxyethyl)phosphine, iodoacetamide, and dithiothreitol, before 3 kDa MWCO filtration. Samples were analyzed using a Waters NanoACQUITY UPLC system with mobile phases consisting of 0.2% FA in H^2^O and 0.2% FA in acetonitrile. For MS/MS analysis of proteins, the NanoACQUITY system was coupled to a Thermo Scientific Orbitrap Fusion Lumos mass spectrometer equipped with the FAIMS Pro interface (19). Proteoform identification was performed with TopPIC (v1.4). Downstream data analysis and quantification was performed using MSstats (v4.0.1), EnhancedVolcano (v1.14.0), and the TopPICR (v0.0.3) R packages.

### Immunoblot Studies

Mouse islets were lysed in a NP-40-based buffer, and analyzed by reducing or non-reducing Tricine-urea-SDS-PAGE (20), and blotted using antibodies listed in Supplementary Table 2, on a LI-COR Odyssey Imaging System. To analyze insulin biosynthesis, islets were pre-incubated in methionine-free RPMI medium for 90 minutes and then treated with L-azidohomoalaine (Invitrogen) and 5 or 25 mM glucose KRB for 90 minutes. Islets were lysed, click-labeled with biotin-alkyne (Invitrogen), immunoprecipitated, and the eluted proteins were analyzed on a Tricine-urea-SDS-PAGE system.

### Immunostaining and Image Analysis

Dispersed mouse islet cells were seeded on chamber slides (Ibidi or Thermo Fisher Scientific) overnight and cultured in 5 or 25 mM glucose RPMI media for the indicated times. For cell proliferation experiments, EdU was added during the last 24 hours of treatment, followed by click-labeling and staining (Invitrogen). TUNEL staining was performed according to the manufacturer’s manual (Roche). For live-cell imaging experiments, cells were labeled with CellRox, MitoSox, TMRM (Thermo Fisher Scientific), and MTG (New England Biolabs), and imaged using a SP5II laser scanning confocal microscope (Leica). Cpd and proinsulin co-stained beta cells were imaged on a SP8X STED white light laser confocal imaging system. Beta cell area were analyzed by immunohistochemistry staining against insulin using a BX61 microscope (Olympus). All antibodies used are listed in Supplementary Table 2. Image analyses were performed using ImageJ (21), QuPath, Ilastik, and CellProfiler pipelines.

### qRT-PCR Experiments

Islet mRNA and DNA were isolated using PureLink RNA Micro (Invitrogen) and QIAamp DNA Micro (Qiagen) kits, and cDNA synthesized using a SuperScript VILO kit (Invitrogen). mRNA and DNA levels were analyzed using SYBR Green-based quantitative real-time PCR (ViiA7 Real-Time PCR System; Applied Biosystems). Primer sequences are listed in Supplementary Table 2.

### Statistical Analysis

Statistical analyses were performed through GraphPad Prism (v9) or R. After conducting normality tests, data sets with normal distribution or with small sample numbers were analyzed using Student’s t-test or ANOVA followed by post-hoc analysis. Data with non-normal distribution were analyzed using the Wilcoxon rank sum test. Statistical significance is indicated in the figures as follows: \**p <* 0.05. All data are presented as mean ± SEM.

### Data and Resource Availability

Data and reagents generated in the current study are available from the corresponding author upon reasonable request.

## Results

### Loss of mature insulin granules and elevated plasma proinsulin in β*Cpe*KO mice

CPE is highly expressed in human and mouse islet endocrine cells (**Figure 1A and 1B**). To study the roles of Cpe in beta cells, we generated beta-cell-specific *Cpe* knockout (β*Cpe*KO) mice by crossing *Ins1*^Cre/+^ and *Cpe*^fl/fl^ mice (**Figure 1C-E**). The deletion of *Cpe* in beta cells leads to near-total loss of mature insulin granules (**Figure 1F**), and significantly elevated fasting plasma proinsulin-like immunoreactivity (**Figure 1G and 1H**).

**Figure 1.**
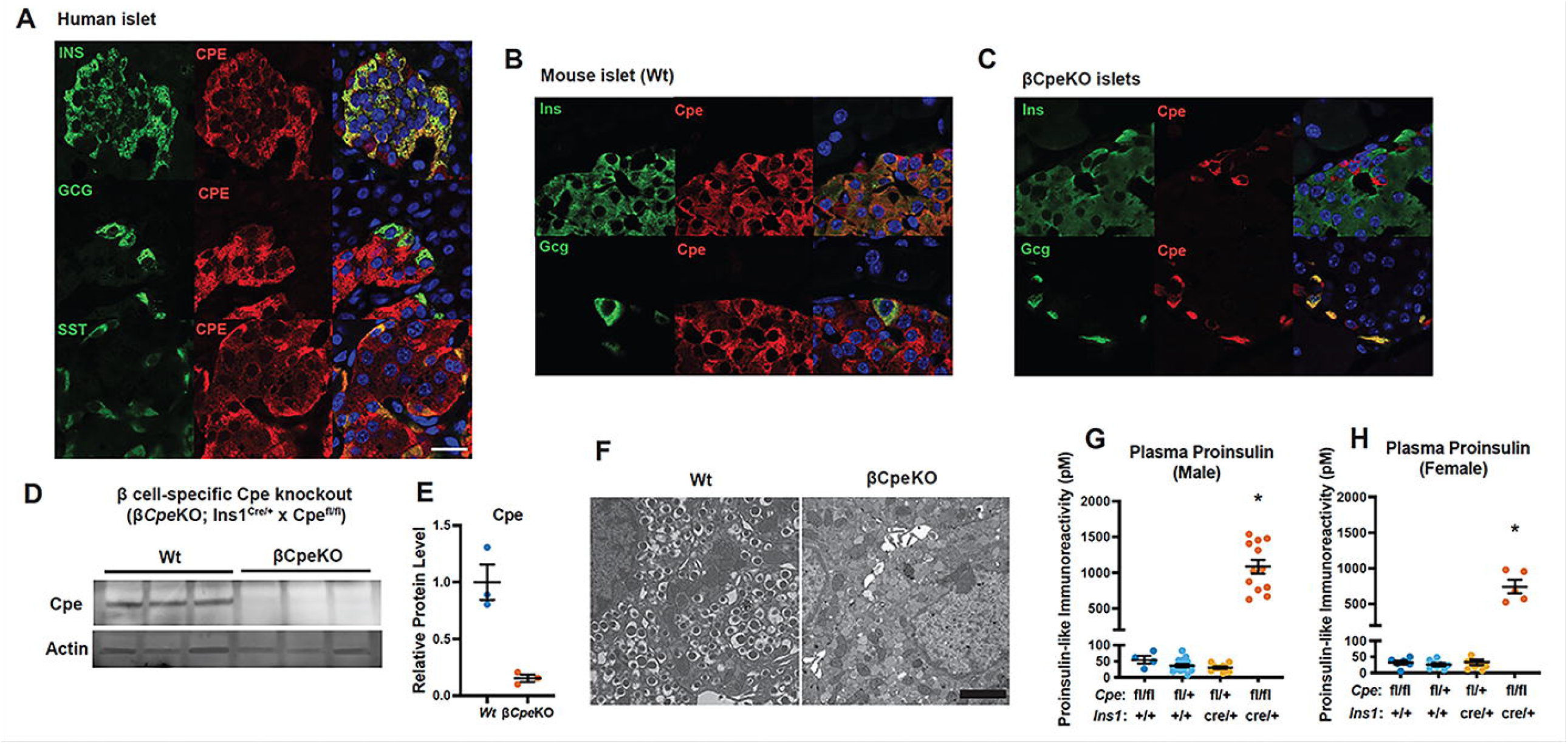
Elevated proinsulin levels in β*Cpe*KO mice. **(A-C)** Expression of CPE in human and mouse pancreatic islet cells was analyzed via immunostaining using antibodies against CPE, insulin (INS), glucagon (GCG), and somatostatin (SST). **(D-E)** Cpe expression in β*Cpe*KO mice was analyzed by immunoblotting (n=3 and 3) using an antibody against Cpe. **(H)** Representative electron micrographs of beta cells from *Wt* and β*Cpe*KO mice. Scale bar=2 µm (n=3 and 3). **(F-G)** 4 hour fasting plasma proinsulin levels in β*Cpe*KO and littermate male and female mice (n>4 per genotype group per sex).

### Permissive peptide processing in β*Cpe*KO islets

Proinsulin is first processed by PC1/3 to form split-32,33 proinsulin with overhanging basic residues. Cpe then removes these basic residues to yield the des-31,32 proinsulin intermediate, which is cleaved by PC2 (or PC1/3) to produce mature insulin following trimming of the remaining basic residues by Cpe (1–3). To study the impact of beta-cell Cpe deficiency on proinsulin processing, we analyzed proinsulin forms using a non-reducing SDS-PAGE system. We found that higher-molecular-weight proinsulin forms were increased in β*Cpe*KO islets (**Figure 2A**). Similar to proinsulin, islet amyloid polypeptide (IAPP) is also synthesized as a larger precursor, proIAPP, and is processed by PC1/3, PC2, Cpe, and Pam to form amidated IAPP (22–24,14). Non-amidated IAPP and intermediate proIAPP (proIAPP^1-48^) forms are increased in β*Cpe*KO islets, although amidated IAPP levels appear comparable between *Wt* and β*Cpe*KO islets (**Figure 2B**). Proteomic assessment confirmed that intact proinsulin levels are increased in β*Cpe*KO islets, while levels of the mature insulin are reduced (**Figure 2C and 2D**). Full-length proIAPP levels are also increased, while levels of amidated mature IAPP level are not reduced in β*Cpe*KO islets (**Figure 2E and 2F**).

**Figure 2.**
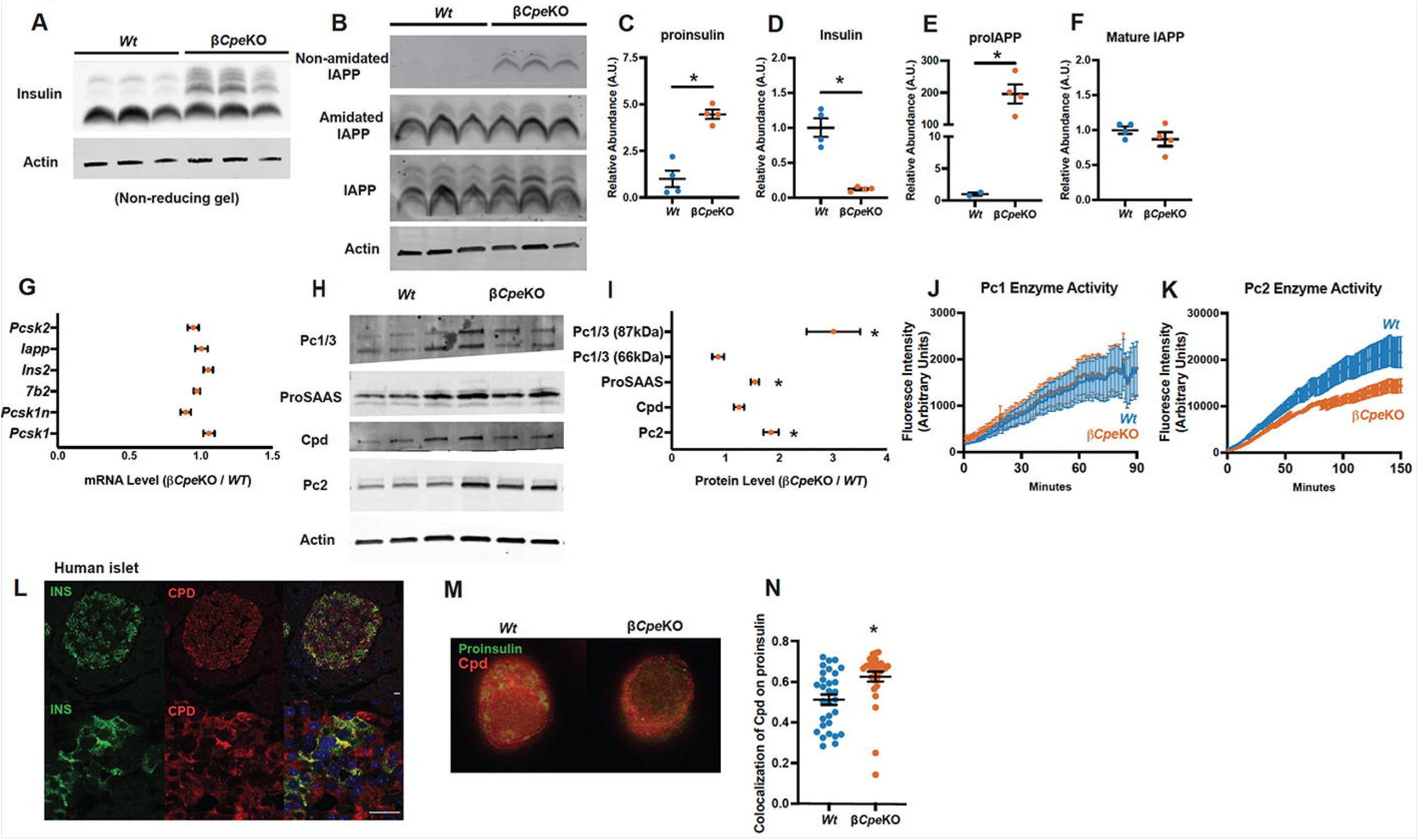
Impaired prohormone processing in β*Cpe*KO islets. **(A)** Islet (pro)insulin levels were analyzed by non-reducing SDS-PAGE and immunoblotting using an antibody against insulin. **(B)** Islet (pro)IAPP levels were analyzed immunoblotting using antibodies against non-amidated IAPP, amidated IAPP, and IAPP. **(C-F)** Quantification of full-length proinsulin, mature insulin B-chain fragment, full-length proIAPP, and amidated mature IAPP, using a top-down proteomics assay by mass spectrometry (n=4 and 4). **(G)** mRNA levels of *Pcsk2, Iapp, Ins2, 7b2, Pcsk1n*, and *Pcsk1* levels in β*Cpe*KO were analyzed by qRT-PCR and presented as folds over *Wt* (n=6 and 6). (**H-I**) Protein levels of proPC1/3 (87kDa), PC1/3 (66kDa), full-length proSAAS, Cpd, and PC2 were analyzed by immunoblotting using antibodies against PC1/3, proSAAS, Cpd, and PC2 (n=3 and 3). (**J-K**) Islet PC1/3 and PC2 enzyme activities were measured by cleavage of synthetic fluorogenic enzyme substrates (n=3 and 3). **(L)** Expression of CPD in human islets beta cells was analyzed by immunostaining using antibodies against Cpd and insulin. Scale bar=25 µm **(M-N)** Immunofluorescence staining of proinsulin and Cpd in *Wt* and β*Cpe*KO beta cells was captured by STED microscope (10 consecutive z-stack images per cell were taken from mice with different genotypes, 3 cells were analyzed per genotype), and colocalization of proinsulin and Cpd was analyzed by Manders method.

To determine whether *Cpe* deletion creates feedback inhibition of peptide hormone maturation, we analyzed prohormone processing enzyme transcript and protein levels. Expression of insulin, IAPP, and processing enzyme transcripts are comparable between β*Cpe*KO and *Wt* islets (**Figure 2G**). Pro-PC1/3 protein (87 kDa) levels are elevated, as are PC2 and proSAAS (**Figure 2H and 2I**); however, total islet PC1/3-specific activity is not changed (**Figure 2J**). PC2-specific activity is reduced in β*Cpe*KO mice (**Figure 2K**), which may occur through increased 7B2-mediated inhibition of PC2 activity (25). We also found that Cpe is not the only carboxypeptidase capable of processing peptide hormones in beta cells. Despite near complete recombination and deletion of *Cpe* (**Figure 1C-1E**), mature insulin remains detectable, and mature IAPP is expressed at similar levels compared to *Wt* islets (**Figure 2B**). Carboxypeptidase D (CPD) has been detected in rodent islet cells and may be sorted to secretory granules (26,27). CPD is expressed in human islet beta cells (**Figure 2L**) and is more restricted to granule-like puncta near cell periphery, as shown in STED super-resolution microscope images (**Figure 2M**). Increased trafficking of Cpd to insulin granules may aid the processing of prohormones in the absence of Cpe, as Manders’ colocalization coefficient of proinsulin and Cpd is significantly higher in Cpe-deficient beta cells (**Figure 2N**).

### β*Cpe*KO mice do not to develop diet-induced obesity and diabetes

Unlike *Cpe* whole body knockout mice or *Cpe* mutant mice (12,13), 8 week-old β*Cpe*KO mice do not develop early-onset obesity and diabetes (males, **Figure 3A-C**; females, **Figure 3D-F**). Islets from β*Cpe*KO mice displayed comparable insulin secretion dynamics to *Wt* islets (**Figure 3G**), speaking against a role for Cpe as a granule sorting receptor in beta cells. Interestingly, proinsulin was released upon high glucose- and KCl-stimulation (**Figure 3H**), suggesting that the proinsulin-containing granules are likely equipped with appropriate granule contents that allow for efficient granule release. Rates of exocytosis were also comparable between *Wt* and *Cpe*-deficient beta cells (**Figure 3I**), although exocytosis events proximal to the plasma membrane, measured by a time-train depolarization experiment, were slightly reduced in beta cells from β*Cpe*KO mice (**Figure 3J**).

**Figure 3.**
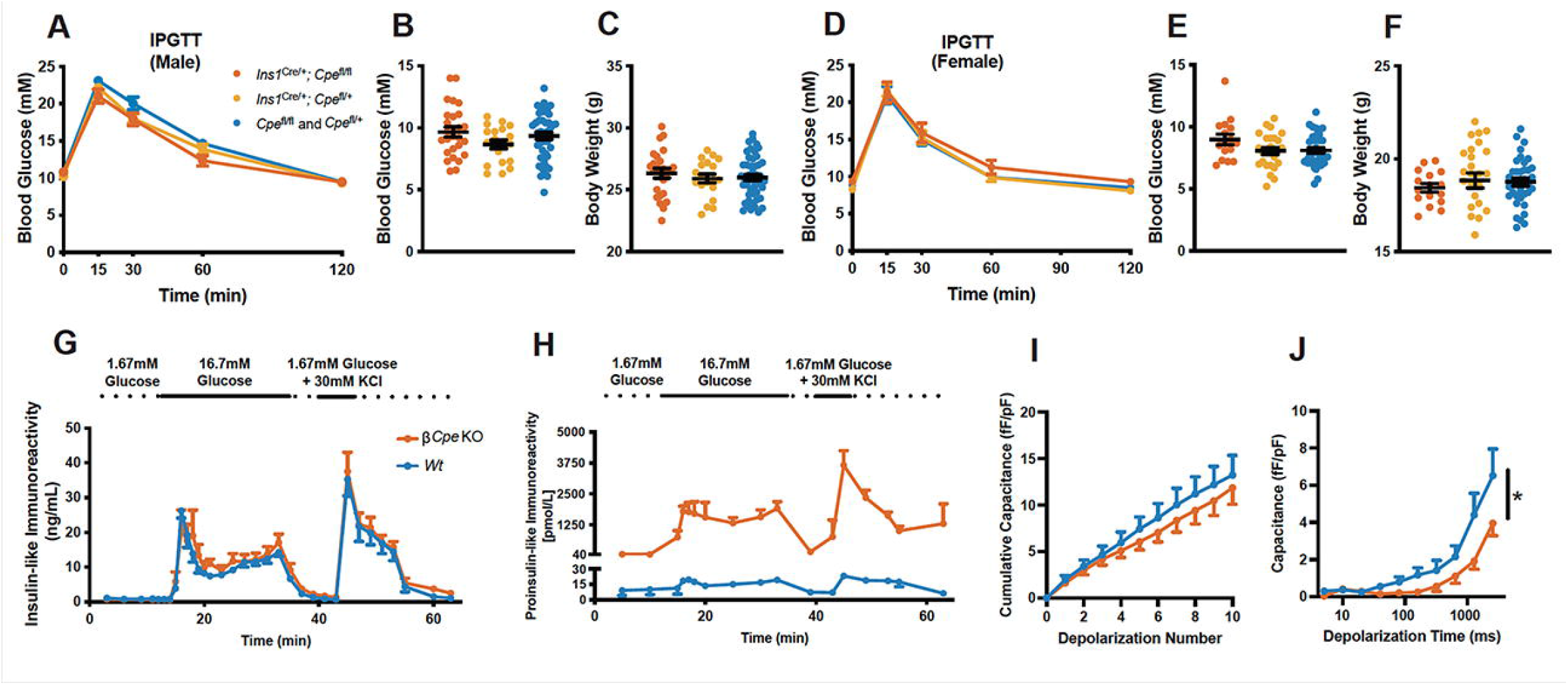
Glucose tolerance and insulin secretion of β*Cpe*KO mice. **(A)** Intraperitoneal glucose tolerance test (IPGTT) was performed on 8 week-old chow-fed β*Cpe*KO and littermate males (n≥5 per genotype). **(B-C)** 4-hour fasting blood glucose levels and body weight of 8 week-old chow-fed β*Cpe*KO and littermate males (n≥5 per genotype). **(D)** IPGTT was performed on 8 week-old chow-fed β*Cpe*KO and littermate females (n≥6 per genotype). **(E-F)** 4-hour fasting blood glucose and body weight of 8 week-old chow-fed β*Cpe*KO and littermate females (n≥6 per genotype). **(G-H)** Insulin-like immunoreactivity and proinsulin-like immunoreactivity during perifusion of 1.67mM glucose, 16.7mM glucose, and 1.67mM glucose plus 30mM KCl (n=5 and 5). **(I-J)** Exocytosis, and exocytosis during a train of depolarization pulses with increased duration, of beta cells from islets of β*Cpe*KO and *Wt* mice (>10 cells per mouse, 3 mice per genotype).

To promote the development of obesity and insulin resistance, we placed 8-week-old β*Cpe*KO mice on a control low-fat diet (LFD, 10% fat) or high-fat diet (HFD, 40% fat), for a duration of 6 months. In the LFD-treated group, β*Cpe*KO mice gained similar weight compared to their *Wt* littermates (males, **Figure 4A**; females, **Figure 4F**), suggesting that the lack of *Cpe* in beta cells does not lead to spontaneous development of obesity. At 20-week post-LFD, male β*Cpe*KO mice displayed modestly increased fasting blood glucose levels (**Figure 4B**); however, their glucose tolerance, insulin tolerance, and percent fat mass, measured at 16 week- and 20 week-post LFD, remained comparable to *Wt* littermates (**Figure 4C-E**). Female β*Cpe*KO mice have slightly elevated fasting blood glucose levels at 4- and 22-week post-LFD, yet displayed similar glucose tolerance, insulin tolerance, and body fat mass compared to littermates (**Figure 4G-J**). Inappropriate proinsulin processing associated with beta-cell *Cpe* deficiency does not contribute to accelerated development of HFD-induced obesity (males, **Figure 4K**; females, **Figure 4P**). Male and female β*Cpe*KO mice showed similar fasting blood glucose levels, glucose tolerance, insulin tolerance, and percent fat mass compared to their littermates (**Figure 4L-O and 4Q-T**).

**Figure 4.**
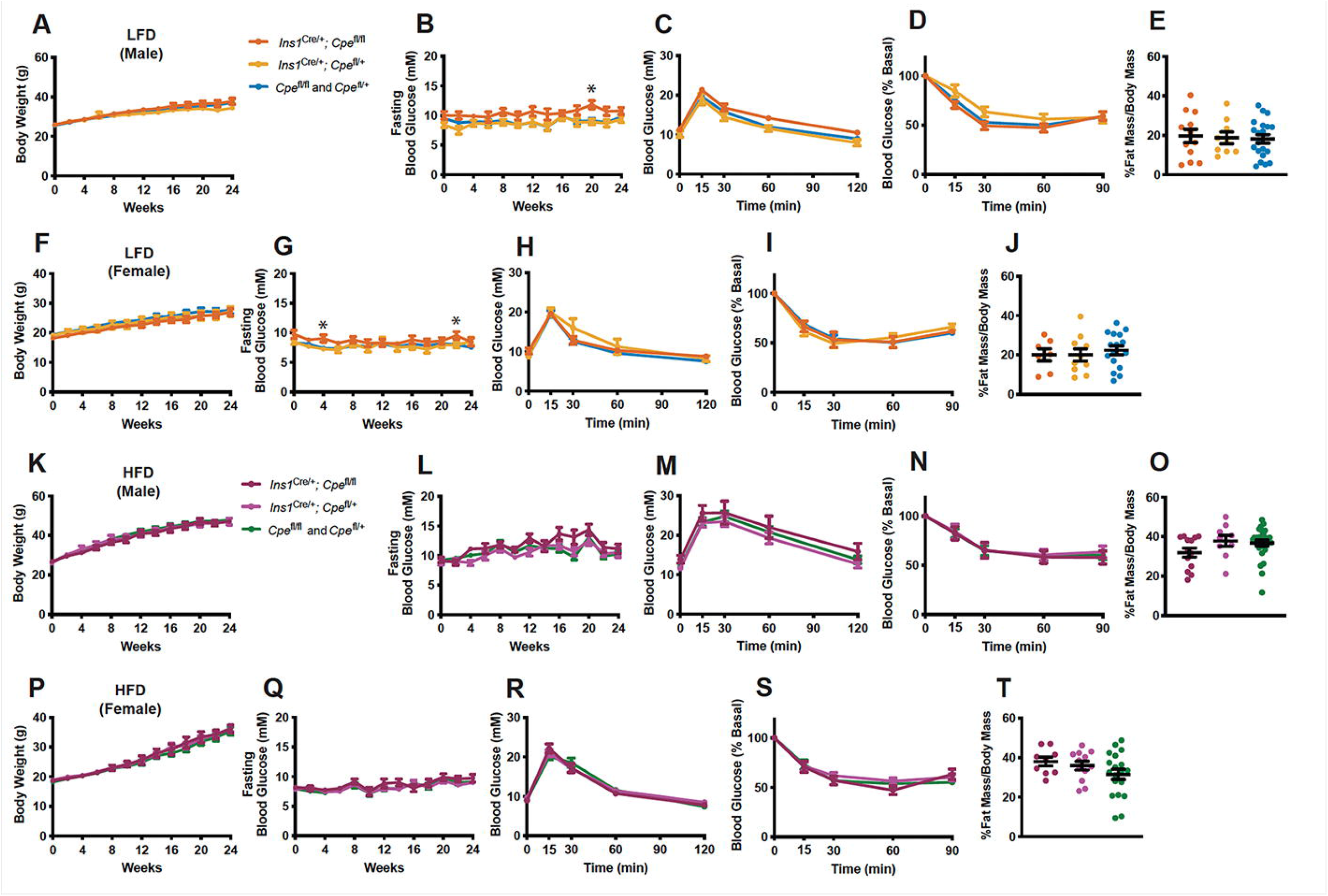
Comparable glucose tolerance and weight gain in HFD-fed β*Cpe*KO and littermate mice. Body weight and 4-hour fasting blood glucose were monitored every 2 weeks for 24 weeks, on low-fat diet (LFD)-fed β*Cpe*KO and their littermate male **(A-B)** and female **(F-G)** mice. IPGTT was performed after 16 weeks of LFD treatment on male **(C)** and female **(H)** mice. Insulin tolerance test (ITT) and body mass composition analysis were performed after 20 weeks of LFD treatment on male **(D-E)** and female **(I-J)** mice (n≥8 per group). Body weight and 4-hour fasting blood glucose were monitored every 2 weeks for 24 weeks, on high-fat diet (HFD)-fed β*Cpe*KO and their littermate male **(K-L)** and female **(P-Q)** mice. IPGTT was performed after 16 weeks of HFD treatment on male **(M)** and female **(R)** mice. ITT and body mass composition analysis were performed after 20 weeks of HFD treatment on male **(N-O)** and female **(S-T)** mice (n≥8 per group).

### Increased beta-cell proliferation in *Cpe*-deficient mice

Despite displaying comparable glucose tolerance, beta-cell area in LFD-treated β*Cpe*KO male and female mice was elevated (**Figure 5A and B**). Although HFD promoted compensatory beta-cell expansion in *Wt* mice, β*Cpe*KO mice failed to increase beta-cell area (**Figure 5C and D**). To study the cause of increased beta-cell area in β*Cpe*KO mice on LFD, we first examined the beta-cell proliferation rate by analyzing the frequency of Ki67^+^ beta cells. However, the number of proliferating beta cells in 6 month-LFD-treated mice was too low to allow for appropriate comparison between β*Cpe*KO and *Wt* mice (data not shown). We therefore isolated islets from 10-week-old mice, cultured them in 5 or 20 mM glucose media for 72 hours, and analyzed EdU incorporation in insulin^+^ beta cells. Beta-cell proliferation was significantly elevated in islets from β*Cpe*KO mice (**Figure 5E**). We also generated inducible *Cpe* knockout mice (iβ*Cpe*KO) by crossing *Pdx1-Cre*^ER^ mice with *Cpe*^flox/flox^ mice (**Figure 5F**). Shortly after oral tamoxifen administration, male, but not female iβ*Cpe*KO mice, become mildly glucose intolerant (**Figure 5G and H**). To induce beta-cell proliferation, we treated iβ*Cpe*KO and their *Wt* littermates with a 60% fat diet for 2 days and analyzed EdU incorporation in the mouse pancreas. We found that the beta-cell proliferation rate was significantly elevated in iβ*Cpe*KO female mice (**Figure 5I**, males; **Figure 5J**, females).

**Figure 5.**
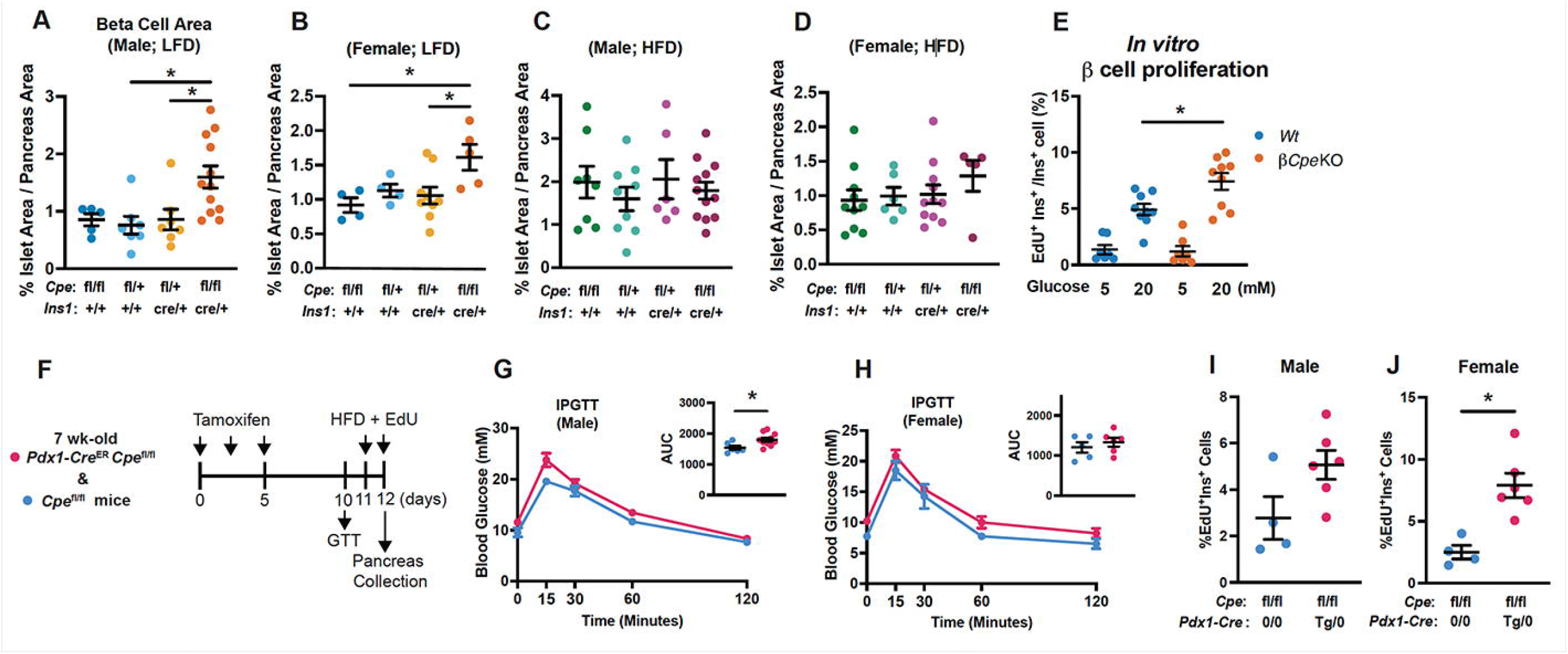
β*Cpe*KO mice have increased beta cell area and beta cell replication. **(A-D)** Beta cell area was analyzed by insulin immunostaining of pancreatic sections from 24 weeks LFD- or HFD-treated male or female β*Cpe*KO and littermate mice (n≥4 per group). **(E)** Beta cell replication rate was analyzed by calculating EdU- and insulin-positive cells in dispersed β*Cpe*KO and *Wt* islets treated with 5mM glucose or 20mM glucose for 72 hours (n≥7 per group). **(F)** Timeline of *in vivo* beta cell replication experiment. **(G-H)** IPGTT was performed in inducible beta cell-specific Cpe knockout mice 5 days after last tamoxifen gavage (n≥5 per group). **(I-J)** Beta cell replication rate was analyzed by calculating EdU- and insulin-positive cells in pancreatic sections from 48-hour 60% HFD-fed β*Cpe*KO and *Wt* male or female mice (n≥4 per group).

### Altered glycolytic gene expression and increased (pro)insulin biosynthesis in *Cpe*-deficient beta cells

To identify the underlying molecular mechanisms contributing to increased beta-cell proliferation in β*Cpe*KO mice, we performed transcriptomic analysis in sorted beta cells from *Wt* and β*Cpe*KO mice (**Figure 6A**). As expected, *Cpe* was drastically reduced in beta cells from β*Cpe*KO mice. Expression of many Hif1alpha-regulated genes (including *Ldha, Hmox1, P4ha1, Pgk1, Mif, Ak4, Bnip3, Pfkp, P4ha2, Slc2a1*, and *Pfkfb3*) was increased. Gene ontology analysis showed that expression of genes related to glycolysis and hypoxia are enriched, while hallmarks of pancreatic beta cells are reduced in *Cpe*-deficient mouse beta cells (**Figure 6B**). We found that the rate of glucose uptake into islet cells is comparable in β*Cpe*KO and *Wt* mice (**Figure 6C**), and glycolytic flux analysis showed that β*Cpe*KO islet cells have a similar oxygen consumption rate compared to *Wt* islet cells (**Figure 6D**). Interestingly, despite having elevated (pro)insulin production (**Figure 6E**) and increased accumulation of proinsulin oligomers (**Figure 6F**), transcripts of canonical unfolded protein response elements were not elevated in β*Cpe*KO beta cells.

**Figure 6.**
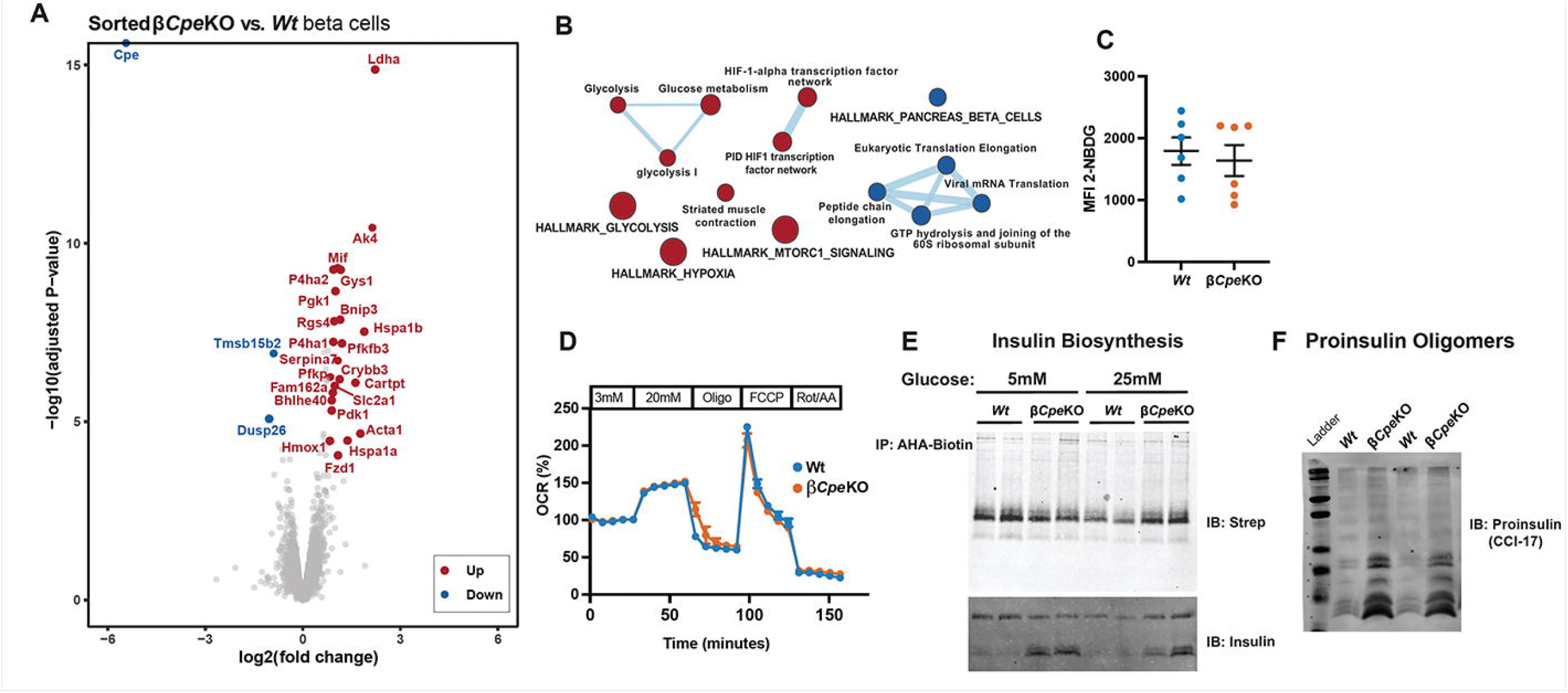
Altered glycolytic transcripts and increased proinsulin biosynthesis in β*Cpe*KO islets. **(A)** Transcriptomic analysis of islet beta cells from 16 week-old chow-fed β*Cpe*KO and *Wt* mice (n=3 and 3). Data were presented as a volcano plot with significantly up- or down-regulated genes annotated. **(B)** Results from Gene Set Enrichment Analysis were presented as EnrichmentMap. Red color: up-regulated gene sets; blue color: down-regulated gene sets. **(C)** Glucose uptake was analyzed by quantifying 2-NBDG fluorescence intensity of dispersed live islet cells from β*Cpe*KO and *Wt* mice via flow cytometer (n=6 and 6). **(D)** Baseline-normalized oxygen consumption rate (OCR) of dispersed islet cells from β*Cpe*KO and *Wt* mice was analyzed (technical triplicate per sample, 3 samples per genotype). **(E)** Freshly isolated islets were equilibrated in methionine-free media, and pulsed with 5mM or 25mM L-azidohomoalaine (AHA) containing media for 90 minutes. Islet protein pellets were click-conjugated with biotin-alkyne, immunoprecipitated with avidin, and analyzed by immunoblotting using streptavidin and an anti-insulin antibody. **(F)** Proinsulin oligomers were analyzed by immunoblotting using an antibody against proinsulin oligomers (mAb CCI-17).

### Dysregulated mitochondrial dynamics and loss of beta-cell identity in glucose challenged β*Cpe*KO islets

Because (pro)insulin production is elevated in β*Cpe*KO islets, we asked whether the morphology or function of the fuel-providing mitochondria is altered in *Cpe*-deficient beta cells. Electron micrograph analysis of mitochondrial images showed that β*Cpe*KO beta cells have similar area, yet reduced size (**Figure 7A-C**). Additional confocal image analysis showed mitochondrial number, area, perimeter, and branch number were all reduced in β*Cpe*KO beta cells (**Figure 7D-I**); while mtDNA content was not reduced (**Figure 7J**). This suggests mitochondria in *Cpe*-deficient beta cells work to accommodate an increased demand for (pro)insulin synthesis. Nevertheless, after prolonged glucose treatment, beta cells from β*Cpe*KO mice had reduced mitochondrial membrane potential upon high glucose stimulation (**Figure 7K**) and displayed elevated levels of mitochondrial and cellular reactive oxygen species (**Figure 7L and 7M**). Beta cells from β*Cpe*KO mice have no increase in mitochondria biogenesis upon high glucose culture, as *Pgc1a* transcript levels are not significantly elevated (**Figure 7N**). Islets from β*Cpe*KO mice failed to display elevated *MafA* transcript level upon high glucose treatment (**Figure 7O**). Rather, *Aldh1a3* transcript levels were significantly elevated, suggesting loss of beta-cell identity in β*Cpe*KO islets (**Figure 7P**). Glucose metabolism was likely altered, as transcript levels of *Pfkp* was significantly increased in β*Cpe*KO islets (**Figure 7Q**). Of note, treatment of islets with high glucose led to increased expression of ER stress markers such as spliced *Xbp1* (*Xbp1s*), but β*Cpe*KO islets showed no increase in *Xbp1s* (**Figure 7R**), inferring that elevated proinsulin biosynthesis does not contribute to increased ER stress in *Cpe*-deficient beta cells.

**Figure 7.**
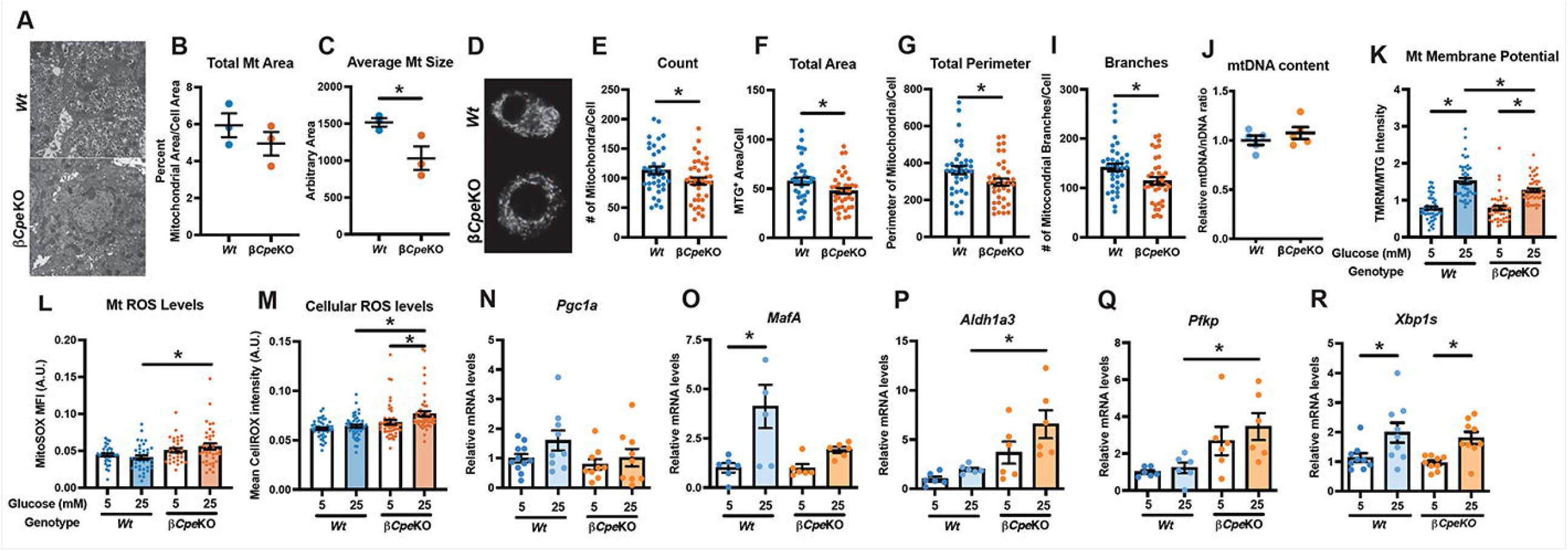
Mitochondrial morphology and function were disturbed in islet cells from β*Cpe*KO mice. **(A)** Representative electron micrographs of mitochondria in beta cells from β*Cpe*KO and *Wt* mice. **(B-C)** Total mitochondria numbers and average mitochondrial size in beta cells were quantified. Each dot represents the average of 2-6 images from one mouse (>400 mitochondria were analyzed per mouse), n=3 per genotype. **(D)** Representative immunofluorescence staining of mitochondria using MitoTracker Green (MTG), in islet cells from β*Cpe*KO and *Wt* mice. **(E-I)** Mitochondria numbers, area, perimeter, and branch number were quantified. Each dot represents one cell, 10-15 cells per mouse, n≥3 per genotype. **(J)** Mitochondrial DNA content (mtDNA) of islets from β*Cpe*KO and *Wt* mice was analyzed by qRT-PCR using ND1 (mtDNA) and 16S (nuclear DNA, nDNA) primer probes, and presented as mtDNA/nDNA ratio. (n=5 per genotype). **(K-M)** Mitochondrial membrane potential, mitochondrial reactive oxygen species (ROS) levels, and cellular ROS levels were analyzed via live cell imaging in 5mM or 25mM glucose-treated dispersed islet cells using Tetramethylrhodamine methyl ester (TMRM), MTG, MitoSOX, and CellROX dyes. Each dot represents one cell, 10-15 cells per mouse, n≥3 per genotype. **(N-R)** qRT-PCR analysis of *Pgc1α, MafA, Aldh1a3, Pfkp, and Xbp1s* in β*Cpe*KO and *Wt* mouse islets treated with 5mM or 25mM glucose for 48 hours (n=6 per group).

### β*Cpe*KO mice have accelerated development of STZ-induced hyperglycemia

We treated β*Cpe*KO and littermate mice with multiple-low-dose-streptozotocin (MLD-STZ) to induce cell dysfunction in a small portion of beta cells, and to create secretory stress in the remaining cells (**Figure 8A**). β*Cpe*KO mice showed higher blood glucose levels at 10 days after the last STZ treatment, compared to *Cpe* heterozygous (β*Cpe*Het) or *Wt* mice (**Figure 8B-D**). β*Cpe*KO mice did not display increased STZ-induced beta-cell death, because beta-cell area (analyzed at 10 days post-STZ) and number of TUNEL^+^ beta cells (analyzed at 3 days post-STZ) was similar in β*Cpe*KO and *Wt* mice (**Figure 8E and F**). Building on our finding of increased beta-cell area in β*Cpe*KO mice (**Figure 5A)**, we found that buffer-treated β*Cpe*KO mice had a higher percentage of beta cells in their islets (**Figure 8G**). Although MLD-STZ treatment led to a modest increase in the percentage of beta cells in *Wt* islets, it reduced the percentage of beta cells in β*Cpe*KO islets (**Figure 8G**). In agreement with previous reports, we showed that the portion of Glut2^+^ beta cells was reduced after MLD-STZ treatment (**Figure 8H**); however, the number of Glut2^+^ beta cells remained comparable in Wt and β*Cpe*KO mice. We also found an increased percentage of Aldh1a3^+^ beta cells upon MLD-STZ treatment (**Figure 8I**). We further analyzed protein levels by mean fluorescence intensity and showed that insulin and Glut2 expression levels were significantly reduced upon STZ treatment in β*Cpe*KO mice (**Figure 8J-L**), suggesting that beta cells in β*Cpe*KO are more susceptible to secretory stress-induced dysfunction and loss of cell identity.

**Figure 8.**
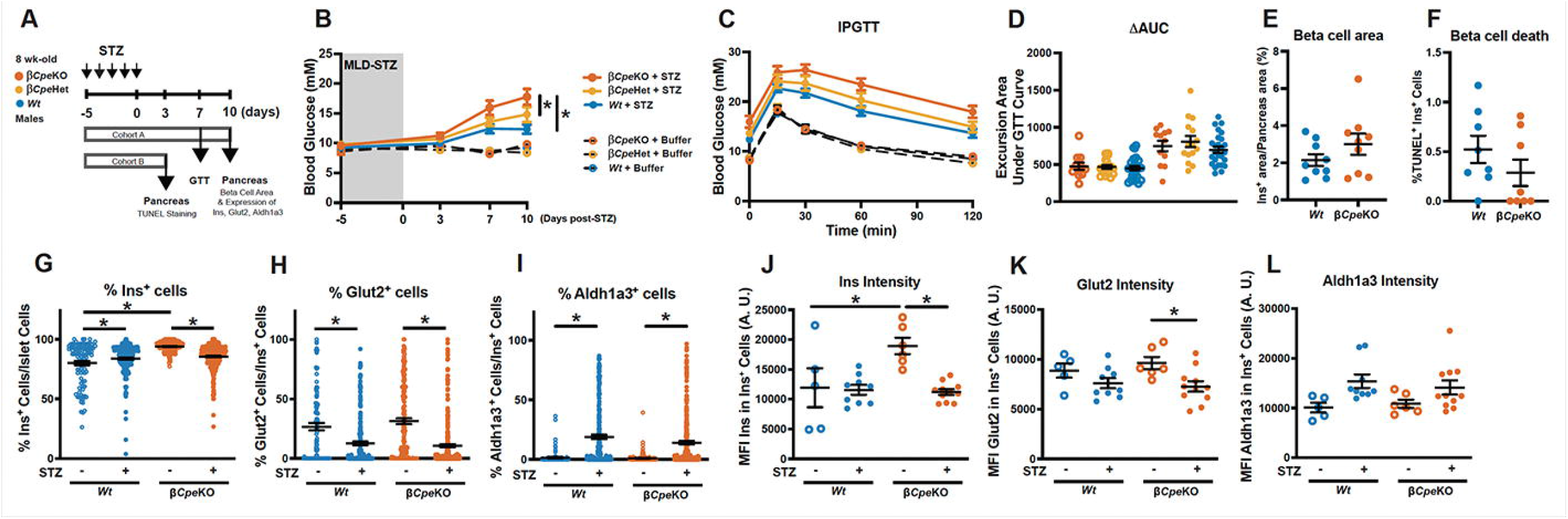
β*Cpe*KO mice have accelerated development of MLD-STZ-induced hyperglycemia. **(A)** Timeline of MLD-STZ experiment. **(B)** Fasting blood glucose levels of β*Cpe*KO and littermate mice treated with a buffer (empty circle with dotted line) or MLD-STZ (filled circle with solid line). **(C-D)** IPGTT was performed 7 days after the last STZ injection, and excursion area under the GTT curve was analyzed (n≥10 per group). **(E)** Beta cell area of MLD-STZ-treated β*Cpe*KO and *Wt* mice (n=9 and 9). **(F)** Beta cell death was analyzed using pancreatic sections of MLD-STZ-treated β*Cpe*KO and *Wt* mice using TUNEL and insulin staining (n=8 and 8). **(G-L)** Frequency and staining intensity of insulin-, glut2-, and aldh1a3-positive cells in pancreatic sections of MLD-STZ-treated β*Cpe*KO and *Wt* mice were analyzed. (**G-I**: each dot represents one islet; **J-L:** each dot represents average fluorescent intensity of islets from one pancreatic section of one mouse; n≥5 per group).

## Discussion

*Cpe* mutations in mice and humans lead to obesity and hyperglycemia; however, the underlying cellular and physiological mechanisms remain unknown. We hypothesized that lack of *Cpe* in pancreatic beta cells is the main contributor to such clinical phenotypes, because: (i) Cpe is required for proper proinsulin processing (3); (ii) reduced expression of Cpe is associated with beta-cell dysfunction in multiple experimental models of diabetes (28–30); and (iii) Cpe may play a protective role in preventing beta-cell death (31). To address this hypothesis, we generated beta-cell-specific *Cpe* knockout mice. Our data indicate that while Cpe is important in normal proinsulin processing, Cpe deficiency alone does not contribute to obesity nor cause marked dysglycemia.

Islets from β*Cpe*KO mice contain markedly more proinsulin peptides, yet have detectable mature insulin peptide, suggesting that another carboxypeptidase, likely Cpd, is compensating. In agreement with an *in vitro* study suggesting that Cpe is essential for PC2-mediated peptide processing (25), we showed that loss of Cpe results in reduced PC2 enzyme activity and increased N-terminally extended proIAPP (which is normally processed by Pc2 into mature IAPP (23)). Although total islet PC1/3 enzyme activity was not changed in β*Cpe*KO mice, PC1/3 protein levels were elevated, suggesting that on a per-enzyme basis, PC1/3 activity is likely reduced in β*Cpe*KO beta cells. To our surprise, even with markedly impaired proinsulin processing and diminished output of mature insulin, β*Cpe*KO mice failed to develop obesity and hyperglycemia spontaneously or when challenged with a high-fat diet; contrary to a recent report showing *Pdx-Cre*^ERT^-mediated *Pcsk1* deletion and elevated proinsulin promote the development of obesity in mice (32). It is plausible that elevated proinsulin, possessing 5% activity compared to insulin (33), is sufficient to maintain glucose homeostasis. Another possibility is that obesity and overt hyperglycemia observed in *Cpe* mutations are driven by insufficient *Cpe* and defective neuropeptide processing in other tissues, such as in the hypothalamus; although mice with *Cpe* deletion in proopiomelanocortin (POMC)-expressing neurons do not become obese (34). It remains to be tested whether Cpe controls body weight and metabolic homeostasis in non-POMC-expressing neuroendocrine cells.

Beta cells are able to adapt to increased insulin demand by increasing production, secretion, and mass prior to hyperglycemia onset (35–37). β*Cpe*KO mice have increased proinsulin production and elevated beta-cell area but remain normoglycemic. Because protein overproduction may change intrinsic metabolic pathways and alter beta-cell fate (38), it is plausible that increased the demand caused by increased insulin production (39) contributes to metabolic pathway rewiring and concomitant beta-cell proliferation in β*Cpe*KO mice. A recent large-scale small molecule screen identified a compound that promotes protein synthesis and beta-cell regeneration. The authors showed that the increased beta-cell regeneration is associated with hypo-translation of mRNAs that are integral to mitochondrial-related processes (40). In support of this idea, we observed altered glycolytic gene signatures, changes in mitochondrial morphology and membrane potential in β*Cpe*KO beta cells. Oxygen consumption rate was not reduced in islets from β*Cpe*KO mice, hinting that additional pathological stimuli are likely needed to disrupt oxidative phosphorylation. Alternatively, metabolic flux analysis may offer more quantitative insights into carbon metabolism and energy flow in islets with inherently elevated proinsulin biosynthesis. It is also possible that the increased proinsulin oxidative folding burden may create ER redox imbalances (41), leading to increased mitochondrial and cellular ROS levels. Future live cell imaging experiments with ROS biosensors could illuminate the cellular sequence of events. Both mild ER stress and ROS have been reported to facilitate beta-cell proliferation (16,42). Despite an accumulation of proinsulin oligomers (43), we failed to detect significant changes in transcripts encoding ER chaperons or unfolded protein response proteins. Of note, beta-cell de-differentiation markers *Serpina7* (44) and *Ldha* (45) are reduced in sorted beta cells from normoglycemic β*Cpe*KO mice, suggesting that the loss of beta-cell identity may occur during early-stage beta-cell compensation prior to the development of hyperglycemia. It is worth mentioning that we have not analyzed possible changes in paracrine signaling in β*Cpe*KO islets.

β*Cpe*KO mice adapted to chronic dietary stress displayed similar weight gain and glucose tolerance as their littermates. We treated β*Cpe*KO mice and their littermates with MLD-STZ to induce acute insulin secretory stress without extensive beta-cell death or loss of beta-cell mass. MLD-STZ treatment led to loss of beta-cell identity in β*Cpe*KO mice, evidenced by an increased percentage of *Aldh1a3*^+^ cells and reduced Glut2^+^ cells in islets. After MLD-STZ treatment, β*Cpe*KO mice also have accelerated development of hyperglycemia, and displayed reduced (pro)insulin and Glut2 expression levels in beta cells. These findings were mirrored *in vitro*, as we observed no induction of *MafA*, and increased *Aldh1a3*, in high glucose-treated β*Cpe*KO islets. We speculate that suboptimal mitochondrial function, or altered cellular redox homeostasis, resulting from the combination of secretory stress and *Cpe* deficiency, contributes to beta-cell dysfunction in MLD-STZ treated *Cpe* deficient islets (46–48). It has been reported that islets from *Cpe* mutant mice are more susceptible to palmitic acid-induced beta-cell apoptosis (31). We were unable to observe detectable differences in TUNEL^+^ beta cells in MLD-STZ treated β*Cpe*KO 3 days after the last STZ injection, when beta-cell apoptosis rates are at their highest (49). Instead, beta cells from β*Cpe*KO mice have reduced Glut2 expression, which may present an adaptive mechanism to attenuate glucose uptake and metabolic-stress induced beta-cell death (50).

In summary, we demonstrated that loss of *Cpe* in pancreatic beta cells does not contribute to spontaneous development of obesity and hyperglycemia in mice. However, elevated proinsulin output likely reshapes beta-cell glucose metabolism and increases its susceptibility to secretory-stress-induced dysfunction and diabetes. Our model may shed light on beta-cell translational adaptation, which likely occurs early during the development of diabetes. Additional studies in other pre-diabetes models and human islets are needed to gain a better understanding of these early adaptive events; and will aid discovery of new therapeutic targets to preserve beta-cell function prior to the onset of diabetes.

## Article information

## Acknowledgments

The authors thank Dr. Paul Orban for helpful comments on the project, Dr. Iris Lindberg for providing the enzyme activity assay protocol and reagents, Dr. Rohit Sharma and Dr. Aaron Cox for providing suggestions on beta cell proliferation experiments, Mr. Daniel Pausula for providing protocol for live cell imaging experiments, Drs. Lei Dei and Galina Soukhatcheva for technical assistance. Mass spectrometry proteomics experiments were performed in the Environmental Molecular Sciences Laboratory, Pacific Northwest National Laboratory, a national scientific user facility sponsored by the DOE under Contract DE-AC05-76RL0 1830.

## Funding

This work is supported by Juvenile Diabetes Research Federation (advanced postdoctoral fellowship 3-APF-2022-1141-A-N to Y-C.C.), Canadian Institutes of Health and Research (grant PJT-153156 to C.B.V.), BC Children’s Hospital Foundation and BC Children’s Hospital Research Institute (Canucks for Kids Fund Childhood Diabetes Laboratories Summer Studentship to K.W.), and National Institutes of Health grants R01DK122160 and U01DK124020 to W-J. Q.

## Duality of Interest

No competing interests relevant to this article were declared.

## Author Contribution

Y-C.C. contributed to study conceptualization, investigation, formal analysis, visualization, and wrote the manuscript; A.J.T contributed to study conceptualization, investigation, and edited the manuscript; J.M.F., X-Q. D., and M.K. contributed to methodology, investigation, and formal analysis; K.W. contributed to investigation, formal analysis, and edited the manuscript; K.F. and A.P. contributed to investigation; A.C.S. contributed to investigation, methodology, and edited the manuscript; R. K-G contributed to methodology and formal analysis; W-J. Q., and P.E.M. contributed to investigation and methodology; C.B.V. contributed to study conceptualization, project administration, resources, reviewed and edited the manuscript. All authors approved the final version of the manuscript. C.B.V. is the guarantor of this work.

## Prior Presentation

Parts of this study were presented at European Association for the Study of Diabetes Annual Meeting in 2018&2022, and Gordon Conference Protein Processing, Trafficking and Secretion Meeting in 2022.

## Notes

### Competing Interest Statement

The authors have declared no competing interest.

