## Supplementary material for "Deletion of carboxypeptidase E in beta cells disrupts proinsulin processing and alters beta cell identity in mice": Sup. Table 1

### Checklist for Reporting Human Islet Preparations Used in Research

Adapted from Hart NJ, Powers AC (2018) Progress, challenges, and suggestions for using human islets to understand islet biology and human diabetes. Diabetologia https://doi.org/10.1007/s00125-018-4772-2

| **Islet preparation** | **1** | **2** | **3** | **4** | **5** | **6** | **7** | **8**[a] |
|---|---|---|---|---|---|---|---|---|
| **MANDATORY INFORMATION** | | | | | | | | |
| Unique identifier | 6030 | R085 | | | | | | |
| Donor age (years) | 30.1 | 61 | | | | | | |
| Donor sex (M/F) | M | M | | | | | | |
| Donor BMI (kg/m$^2$) | 27.1 | 30.5 | | | | | | |
| Donor $HbA_{1c}$ or other measure of blood glucose control | C-peptide 2.54 ng/mL | 5.6 | | | | | | |
| Origin/source of islets[b] | nPOD | ADI IsletCore | | | | | | |
| Islet isolation centre | U of Florida | U of Alberta | | | | | | |
| Donor history of diabetes? Please select yes/no from drop down list | | | | | | | | |
| **If Yes, complete the next two lines if this information is available** | | | | | | | | |
| Diabetes duration (years) | | | | | | | | |
| Glucose-lowering therapy at time of death[c] | | | | | | | | |

*Continues on the next page*

| RECOMMENDED INFORMATION | | |
|---|---|---|
| Donor cause of death | Head Trauma | NDD-Neurological |
| Warm ischaemia time (h) | N/A | N/A |
| Cold ischaemia time (h) | N/A | N/A |
| Estimated purity (%) | N/A | N/A |
| Estimated viability (%) | N/A | N/A |
| Total culture time (h)[d] | N/A | N/A |
| Glucose-stimulated insulin secretion or other functional measurement[e] | N/A | N/A |
| Handpicked to purity? Please select yes/no from drop down list | | |
| Additional notes | Pancreatic sections | Pancreatic sections |

[a]If you have used more than eight islet preparations, please complete additional forms as necessary
[b]For example, IIDP, ECIT, Alberta IsletCore
[c]Please specify the therapy/therapies
[d]Time of islet culture at the isolation centre, during shipment and at the receiving laboratory
[e]Please specify the test and the results
