## Supplementary material for "Deletion of carboxypeptidase E in beta cells disrupts proinsulin processing and alters beta cell identity in mice": Sup. Table 2

**Supplementary Table 2**

| Antibodies: | Sources: | Catalog Number: |
|---|---|---|
| anti-Cpe | Millipore Sigma | #AB5314 |
| anti-Pc1/3 | Abcam | #ab191452 |
| anti-Pc2 | Abcam | #ab135808 |
| anti-ProSAAS | Millipore Sigma | #ABN2268M |
| anti-Cpd | Bethyl Laboratories | #A305-514A |
| anti-insulin | Cell Signaling | #L6B10 |
| anti-insulin | Dako | #IR002 |
| anti-proinsuiln | Abnova | #MAB2868 |
| anti-amidated IAPP | MedImmune | #F025 |
| anti-non-amidated IAPP | MedImmune | #F084 |
| anti-IAPP | Peninsula Laboratories | #T-4145 |
| anti-actin | Abcam | #ab8226 |
| IRDye-conjugated secondary antibodies | LI-COR | #926-68070 and #926-32211 |
| Goat anti Rabbit IgG (H+L) Secondary Antibody, Oregon Green 488 | Invitrogen | #O11038 |
| Biotin-SP-AffiniPure Donkey Anti-Mouse IgG (H+L) | Cedarlane Labs | #715-065-150 |
| v500 Streptavidin | BD | #561419 |
| Alexa 488 goat anti-guinea pig | Thermo Fisher Scientific | #A11073 |
| Alexa 594 goat anti-mouse | Thermo Fisher Scientific | #A32742 |
| Alexa 647 goat anti-rabbit | Thermo Fisher Scientific | #A32733 |
| **Dyes and Probes:** | **Sources:** | **Catalog Number:** |
| IRDye-conjugate Streptavidin | LI-COR | #926-32230 |
| CellRox | Thermo Fisher Scientific | #C10422 |
| MitoSox | Thermo Fisher Scientific | #M36008 |

| | | |
|---|---|---|
| TMRM | Thermo Fisher Scientific | #T668 |
| MTG | New England Biolabs | #9074S |
| TUNEL staining reagents | Roche | #11684795910 |
| Click EdU reagents | Invitrogen | #C10337 |
| 2-NBDG | Thermo Fisher Scientific | #N13195 |
| **Chemicals and Consumables:** | **Sources:** | **Catalog Number:** |
| Streptozotocin | Sigma-Aldrich | #S0130-100MG |
| EdU | Toronto Research Chemicals | #E932175 |
| Biotin-alkyne | Invitrogen | #C33372 |
| L-Azidohomoalanine | Invitrogen | #C10102 |
| High Capacity NeutrAvidin Agarose | Thermo Scientific | #209202 |
| μ-Slide 8 Well | Ibidi | #800826 |
| Lab-Tek™ II Chamber Slide | Thermo Scientific | #154534PK |
| **Commercial Assays:** | **Sources:** | **Catalog Number:** |
| Rodent insulin chemiluminescence ELISA | Alpco | #80-INSMR |
| Rodent proinsulin ELISA | Mercodia | #10-1232-01 |
| Lactate ELISA | Abcam | #ab65330 |
| RNeasy Plus Micro kit | Qiagen | #74034 |
| QIAamp DNA Micro kit | Qiagen | #56304 |
| PureLink RNA Micro kit | Invitrogen | #12183016 |
| SuperScript VILO kit | Invitrogen | #11754050 |
| Fast SYBR Green Master Mix | Applied Biosystems | #4385612 |
| **Rodent Diets:** | **Sources:** | **Catalog Number:** |
| Chow | Teklad | #2918 |
| Low fat diet (10% fat) | Research Diets | #D12450H |
| High fat diet (45% fat) | Research Diets | #D12451 |
| High fat diet (60% fat) | Research Diets | #D12492 |

| **Primers:** | **Forward Sequence** | **Reverse Sequence** |
|---|---|---|
| *Rplp* | TCGGGTCCTAGACCAGTGTTC | AGATTCGGGATATGCTGTTGGC |
| *Cpe* | CAGCAAGAGGACGGCATCTC | GTCCAACCGCCTCATTACCAT |
| *Pcsk1* | CTTTCGCCTTCTTTTGCGTTT | TCCGCCGCCCATTCATTAAC |
| *Pcsk2* | GTGTGATGGTTTTTGCGTCTG | GGGAGCTTTCGGACTCCAA |
| *proSAAS* | CGACGAGACTCCTGACGTG | GCACCTCGGGACCCAAATC |
| *Ins2* | GCTTCTTCTACACACCCATGTC | AGCACTGATCTACAATGCCAC |
| *Iapp* | CGGACCACTGAAAGGGATCTT | CGTGTTGCACTTCCGTTTGT |
| *7b2* | CCTCAAGGCTGGTCTCTGCTA | GGCCCAACAAGATTCATGGC |
| *Nd1* | CTAGCAGAAACAAACCGGGC | CCGGCTGCGTATTCTACGTT |
| *16s* | CCGCAAGGGAAAGATGAAAGAC | TCGTTTGGTTTCGGGGTTTC |
| *Aldh1a3* | TTCTTGGGCATGTGCTTAGA | AGCCTGAAGCCCAAATACCAA |
| *Pgc1a* | TATGGAGTGACATAGAGTGTGCT | CCACTTCAATCCACCCAGAAAG |
| *MafA* | AGGAGGAGGTCATCCGACTG | CTTCTCGCTCTCCGAAATGTG |
| *Pfkp* | CTTGGGACAAAACGCACCCT | GGAGACAGTAGCAGGAACCAT |
| *Xbp1s* | GGTCTGCTGAGTCCGCAGCAGG | AGGCTTGGTGTATACATGG |
